# tRNA dissociation from EF-Tu after GTP hydrolysis and P_i_ release: primary steps and antibiotic inhibition

**DOI:** 10.1101/602383

**Authors:** Malte Warias, Helmut Grubmüller, Lars V. Bock

## Abstract

In each round of ribosomal translation, the translational GTPase EF-Tu delivers a tRNA to the ribosome. After successful decoding, EF-Tu hydrolyses GTP, which triggers a conformational change that ultimately results in the release of the tRNA from EF-Tu. To identify the primary steps of these conformational changes and how they are prevented by the antibiotic kirromycin, we employed all-atom explicit-solvent Molecular Dynamics simulations of the full ribosome-EF-Tu complex. Our results suggest that after GTP hydrolysis and Pi release, the loss of interactions between the nucleotide and the switch 1 loop of EF-Tu allows domain D1 of EF-Tu to rotate relative to domains D2 and D3 and leads to an increased flexibility of the switch 1 loop. This rotation induces a closing of the D1-D3 interface and an opening of the D1-D2 interface. We propose that the opening of the D1-D2 interface, which binds the CCA-tail of the tRNA, weakens the crucial EF-Tu-tRNA interactions which lowers tRNA binding affinity, representing the first step of tRNA release. Kirromycin binds within the D1-D3 interface, sterically blocking its closure, but does not prevent hydrolysis. The resulting increased flexibility of switch 1 explains why it is not resolved in kirromycin-bound structures.

## Introduction

Elongation factor Tu (EF-Tu) is a central part of the bacterial translation machinery. During each round of translation elongation, EF-Tu delivers an aminoacyl-tRNA (aa-tRNA) to the ribosome in a ternary complex with GTP [1] (Fig. 1a). The successful decoding of the mRNA codon by the aa-tRNA leads to a closing of the small ribosomal subunit (30S), which in turn docks EF-Tu at the sarcin-ricin loop of the large subunit (50S) in the GTPase-activated state [2, 3]. Subsequently, GTP is hydrolyzed to GDP, followed by the release of inorganic phosphate (P_i_) and a conformational change of EF-Tu [4]. Because the conformational change of EF-Tu occurs rapidly and its rate is limited by P_i_ release, EF-Tu was suggested to behave like a “loaded spring” whose tension is relaxed after P_i_ dissociation [4]. This conformational change eventually leads to the tRNA being released from EF-Tu, followed by the accommodation of the tRNA into the A site and the dissociation of EF-Tu from the ribosome [5]. Ensemble and single-molecule kinetic experiments indicate that EF-Tu dissociates, first, from the 3’ end of the tRNA and from the GTPase associated center (GAC) of the ribosome and subsequently from the rest of the tRNA [6]. It has been suggested that tRNA accommodation proceeds in a step-wise manner [7, 8] and that EF-Tu helps the tRNA to assume an intermediate step before full accommodation [9].

**Figure 1:**
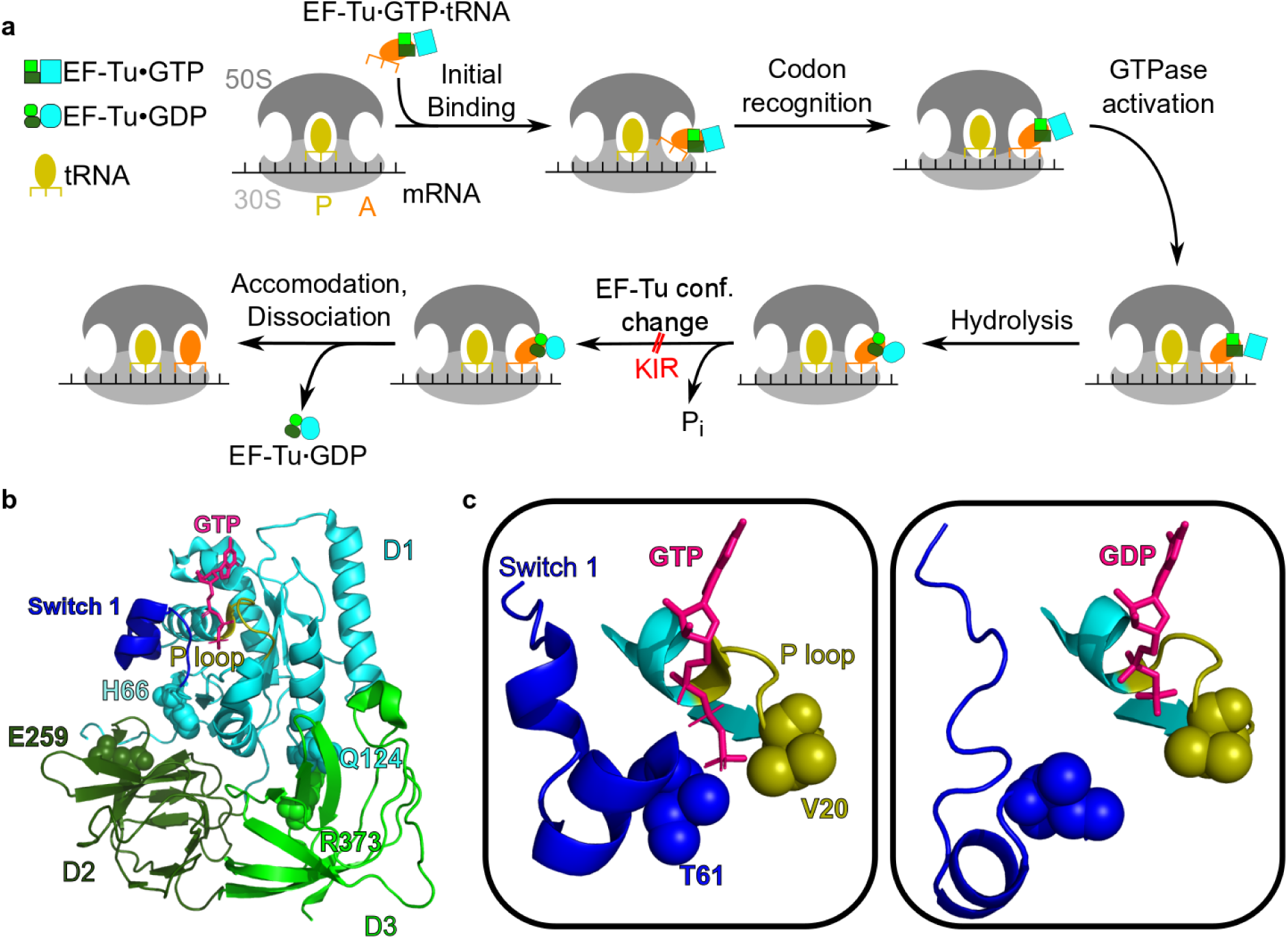
(**a**) Sketch of the mechanism by which EF-Tu delivers aa-tRNA to the ribosomal A site. EF-Tu is represented by three boxes (light green, dark green, cyan) and the aa-tRNA is shown in orange. (**b**) Cartoon representation of EF-Tu in the GTPase-activated conformation [3]. Residues discussed in the text are shown as spheres and GTP as sticks. (**c**) Switch 1 and P loop shown as cartoon representations for the GTPase-activated conformation and for a simulation snapshot containing GDP. The residues closest to the *γ*-phosphate of GTP are shown as spheres.

EF-Tu consists of the GTP-binding domain D1 and two *β*-barrel domains D2 and D3 [10] (Fig. 1b). All three domains bind the tRNA: the CCA-tail of the tRNA along with the attached amino acid is bound to a cleft between D1 and D2, while D3 interacts with the T stem of the tRNA [11]. A closed and an open conformation of the free ternary complex have been solved in complex with GTP and GDP, respectively [10, 12]. The main difference between the two conformations is a 100° rotation of D1 against D2 and D3 and a different fold of the switch 1 region which is a part of D1. EF-Tu bound to the ribosome with non-hydrolyzable GTP [3, 13] assumes a similar conformation as the isolated ternary complex [11]. The switch 1 region was observed in an *α*-helical conformation for closed GTP-bound EF-Tu [10] and in a *β*-sheet conformation for open GDP-bound EF-Tu [14]. These large conformational changes between GTP- and GDP-bound EF-Tu have been suggested to induce the release of the tRNA from EF-Tu and the dissociation of EF-Tu from the ribosome [15]. Recently, however, a crystal structure of EF-Tu with a non-hydrolysable GTP analogue (GDPNP) has been resolved in an open conformation which together with smFRET experiments suggests that, when free in solution, GTP-bound EF-Tu samples a wide range of conformations [16]. These results challenge the model that the identity of the nucleotide, GTP or GDP, acts as a switch for the conformations of EF-Tu. Further, smFRET experiments using dyes attached to different locations on EF-Tu suggest that GTP hydrolysis results in a smaller conformational change of ribosome-bound EF-Tu that is not compatible with the full transition to the open conformation [17]. In the GTPase-activated conformation of ribosome-bound EF-Tu, before GTP hydrolysis, the *γ*-phosphate of GTP interacts with EF-Tu via the P-loop (V20, D21), the switch 1 loop (T61) and the switch 2 loop (G83) [18, 3, 13]. The switch 1 loop in turn is involved in the binding of EF-Tu to the tRNA (nucleotides 1–3 and 73–75). The conformation of EF-Tu after GTP hydrolysis and P_i_ release and before dissociation in the absence of antibiotics has not been structurally resolved yet.

Kirromycin (KIR) is an antibiotic that directly binds to the interface of EF-Tu domains D1 and D3 and prevents dissociation of EF-Tu from the ribosome and from the aa-tRNA after GTP hydrolysis [19, 20, 21, 15, 22]. Structures of ribosome-bound EF-Tu with GDP and KIR have been obtained by x-ray crystallography and cryo-EM [15, 22]. With KIR bound, the overall conformation of EF-Tu remains close to the GTP-bound conformation after hydrolysis, both on and off the ribosome [20, 23, 24, 25, 26, 27, 21, 15, 22]. However, there seems to be a conformational difference between ribosome-bound EF-Tu in complex with GTP and with GDP+KIR. In the ribosome-bound structures of either wt EF-Tu with non-hydrolysable GTP analogues [18, 3, 13] or of a hydrolysis-incompetent EF-Tu variant (H84A) with GTP [13], the switch 1 loop is in the *α*-conformation. In the high-resolution structures with KIR+GDP [21, 15, 22], however, the switch 1 loop is not resolved, suggesting that it is more flexible in this state.

To resolve these experimental discrepancies and to elucidate how the antibiotic KIR prevents both dissociation of the tRNA from the ternary complex and release of the complex from the ribosome, we have carried out atomistic simulations of the full ribosome in explicit solvent. We investigated the conformational changes of the ternary complex after GTP hydrolysis and P_i_ release while bound to the ribosome in the absence of KIR. Further, we addressed the question of how these changes might lead to dissociation of the tRNA from EF-Tu and how they are prevented by KIR. To that aim, we used all-atom explicit-solvent molecular dynamics (MD) simulations started from a cryo-EM structure of EF-Tu in complex with GDP and Phe-tRNA^Phe^ bound to the *E. coli* ribosome [22] with and after removal of KIR. In the absence of KIR, we found intermediate conformations of EF-Tu resulting in a weakended interaction between EF-Tu and the amino acid attached to the tRNA, suggesting that this intermediate prepresents a primary step towards the dissociation of the tRNA from EF-Tu and its accommodation into the A site.

## Methods

### Ribosome complex Molecular Dynamics simulations

In order to investigate the conformational changes of the ribosome-EF-Tu complex and the effect of KIR, we carried out all-atom explicit-solvent molecular dynamics simulations in the absence and presence of KIR. As a starting structure, the cryo-EM structure of an *E. coli* ribosome in complex with tRNAs, EF-Tu, GPD, and KIR was used [22]. Residues 41–62 of EF-Tu were not resolved in the cryo-EM structure. To model these residues, a homology model EF-Tu was build using SWISS-MODEL [28] with the sequence of E.Coli EF-Tu and GTP-bound x-ray structure of T. aquaticus EF-Tu (pdb id: 1EFT [29]) as a structural template. After rigid-body fitting of residues 24–40 and 63–71 of the homology model to the corresponding residues in the cryo-EM structure residues 41–62 of the homology model were included in our model. Protein L1 and nucleotides of H38 were also not resolved and added to our model as described previously [30].

For the simulations without KIR, KIR was removed from the starting structure. All simulations were carried out using the GROMACS software package version 5.1 [31] with the amber99sb forcefield [32], and the SPC/E water model [33]. Parameters from Joung and Cheatham [34] and from Aduri et al. [35] were used for K^+^Cl^−^ ions and modified nucleotides, respectively. Initial coordinates of KIR were extracted from the cryo-EM structure [22] and then protonated and energy-minimized by a HF/6-31G* optimization in GAUSSIAN 09 [36]. The electrostatic potential, calculated from the optimized structure at more than 140,000 points of a molecular surface around the KIR, was fitted to partial charges placed at the atomic positions using the ESPGEN module in AMBER 11 [37]. Additional force field parameters for KIR were obtained using the ANTECHAMBER module in AMBER. The parameters were converted for use in GROMACS using ACPYPE [38]. Long-range electrostatics were calculated using the PME method with grid spacing of 0.12 nm and a cut-off of 1 nm [39]. Van-der-Waals interactions were calculated for atoms within 1 nm of each other.

The complex was placed in a dodecahedron box with a minimum distance of 1.5 nm between box borders and any solute atom, and then the box was solvated with water molecules. Next, the charge of the system was neutralized by adding K^+^ ions using the GENION program from the GROMACS package. Then, Mg^2+^and Cl^−^ ions with a concentration of 7 mM and K^+^Cl^−^ ions with a concentration of 150 mM were added.

For each case, with and without KIR, we ran two independent simulations. Each simulation system was equilibrated in four steps as described previously [30]:

- Energy minimization using steepest decent.
- 0–50 ns: Position restraints on all atoms that were resolved in the cryo-EM structure (force constant *k*=1000 kJ/mol/nm^2^) and Berendsen barostat [40] (*τ*_*p*_=1 ps).
- 50–70 ns: Linear decrease of position restraint force constant *k* to zero.
- 70–2070 ns: No position restraints and Parrinello-Rahman barostat [41] (*τ*_*p*_=1 ps).

In all simulations, the bond lengths were constrained using the LINCS algorithm [42]. Virtual site constraints [43] and an integration step of 4 fs were used. The temperature of solute and solvent was controlled independently using velocity rescaling [44] (*τ*_*T*_ =0.1 ps).

Production runs consisted of 2 *µ*s of unrestrained MD simulations and only these trajectories, recorded every 5 ps, were used for analysis.

### Molecular Dynamics Simulations of EF-Tu in solution

To test whether the opened D1-D3 interface is energetically unfavourable, we performed simulations of EF-Tu in complex with aa-tRNA and GTP in solution started from a conformation with an open D1-D3 interface. The starting structure was taken from a GTPase activated ribosome·EF-Tu·aa-tRNA structure [3]. All ions resolved in the cryo-EM structure within a 15-Å radius around the tRNA and EF-Tu were kept for the simulation system. The setup then followed the same steps as the simulations of the ribosome complex (see above).

### Principal component analysis

To obtain the dominant conformational modes of motion for EF-Tu and the tRNA, we employed principal component analysis (PCA) [45]. The eigenvectors of the covariance matrix were calculated using the program gmx covar from the GROMACS package [31]. In order to identify the modes of EF-Tu’s domain 1 relative to the other two domains, we first rigid-body fitted the nitrogen and both carbon atoms of the backbone of residues 210-294 and 303-393 for each frame of each trajectory to the corresponding atoms of the cryo-EM structure [22]. The covariance matrix was then constructed from the concatenated trajectories of nitrogen and carbon atoms of the backbone of residues 11-198 (D1).

### Residue distances

To measure the opening of the domain interfaces of EF-Tu, which bind either the CCA-tail of the tRNA or KIR, distances between EF-Tu residues as a function of simulation time were calculated from the trajectories. For the KIR binding site located between domains D1 and D3, the distance between the C*α* atoms of residues Q124 and E373 was calculated. The opening of the tRNA binding cleft was observed by measuring the distance between the C*α* atoms of residues H66 and E259.

### Volume Overlap

To test whether the conformational motion of D1 relative to D2 and D3 observed in simulations without KIR can also occur in the presence of KIR, we investigated if this motion would lead to a clash between KIR and EF-Tu. To that aim, we first extracted snapshots at an interval of 4 ns from trajectories. These snaphots were then rigid-body fitted to the cryo-EM structure using coordinates of all atoms of D1 of EF-Tu. Next, the overlap of the Van-der-Waals volume from EF-Tu from the trajectory with that of the KIR from the cryo-EM structure was calculated. To calculate this overlap, we first calculated the Van der Waals volumes *V*_EF-Tu_, *V*_KIR_, and *V*_EF-Tu+KIR_ for the EF-Tu atoms, for the KIR atoms, and for the combined set of atoms, respectively, using Monte Carlo integration. To this aim, all EF-Tu atoms within 1 nm of KIR were selected, and then 20 10^6^ points were randomly placed in a box defined by extensions of the atomic coordinates. The volume is then calculated by *V* = (Number of points inside Van-der-Waals sphere)/(Total number of points)*·*(Box volume). The overlap *V*_overlap_ is then gained by *V*_overlap_ = *V*_EF-Tu_ + *V*_KIR_ − *V*_EF-Tu+KIR_.

### Free-energy calculation of D1-D3 interace closure

To estimate the the free-energy difference between EF-Tu with an open and closed D1-D3 interface, we used Umbrella sampling simulations [46] along a reaction coordinate representing the closing motion. To obtain the reaction coordinate, first, the trajectories of the ribosome-EF-Tu complexes, were rigid-body fitted using D1 atoms to the cryo-EM structure, and, second, a PCA of the D3 atom coordinates was performed (atom selection as described in PCA section). The first eigenvector obtained from the PCA was chosen as the reaction coordinate. The first simulation of the isolated ternary complex that showed a transition from the open to the closed conformation (sim5) was projected onto this reaction coordinate. Then, 51 equally spaced projections were chosen as windows for the umbrella sampling. For each window, first a structure corresponding to the projection values was extracted. Next, from each of these structures, a 75-ns MD simulation was started with an additional umbrella potential centered at the corresponding projection value and a spring constant of 100 kJ/mol/nm^2^. The first 10 ns of each trajectory were discarded to allow for equilibration. Free energies were calculated using the g wham tool and the errors were estimated by 300 bootstrapping steps [47].

### Interaction enthalpies

To estimate whether the conformational motion of EF-Tu, observed in the simulations, changes the binding strength between EF-Tu and the aa-tRNA, we calculated interaction enthalpies between EF-Tu and the amino acid bound to the tRNA were calculated for residues in close contacts. The close contacts were identified using the program g contacts [48] with a minimum distance of 3 Å. Interaction enthalpies were calculated every 5 ps, using the rerun using the rerun option of the GROMACS package.

## Results

After GTP hydrolysis and P_i_ release, EF-Tu changes its conformation, ultimately resulting in tRNA accommodation and EF-Tu release. To investigate the conformational changes of EF-Tu after P_i_ release, we performed MD simulations of the ternary complex (EF-Tu·GDP·aa-tRNA) bound to the ribosome. The simulations were started from a cryo-EM structure of the ribosome-bound ternary complex containing KIR [22]. The unresolved switch 1 loop was modeled based of its GTP-bound conformation [29]. Four MD simulations were carried out, each 2 *µ*s in length: two simulations including and two simulations after the removal of KIR.

### Dynamics of switch 1 loop after P_i_ release

In the GTP-bound conformation, switch 1, switch 2 and P-loop residues of EF-Tu are in contact with GTP [18, 3, 13]. The only direct contact of the switch 1 loop with GTP is between Thr61 and the *γ*-phosphate [18] (Fig. 1c) This contact is lost after hydrolysis and P_i_ release. To test if the loss of this contact results in either different contacts or in a complete loss of contacts between the switch 1 region and the nucleotide, we monitored the contact occupancies of the *β*-phosphate with the P-loop and switch 1 region over the course of our simulations, which included GDP and were started from the GTP-bound form of the switch 1 region (Fig. 2a).

**Figure 2:**
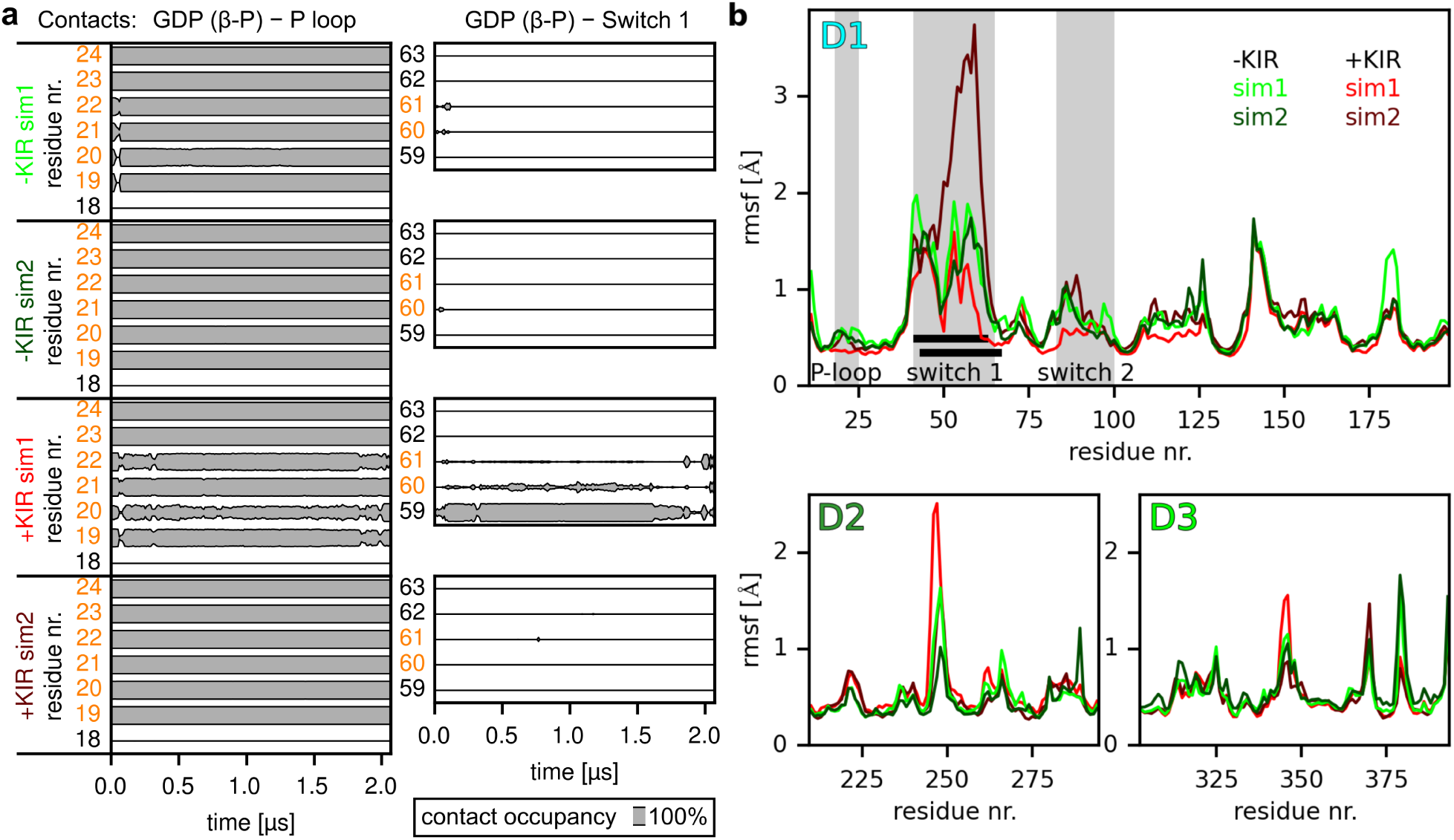
(**a**) Contacts of EF-Tu residues with the *β*-phosphate of GDP or GTP. For residues of the P-loop of EF-Tu (left panel) and the switch 1 loop (right panel), the occupancy of contacts with the *β*-phosphate of GDP (distance below 5 Å) was monitored over the course of the simulations with and without KIR (four rows). Residues that are in contact with the *β*-phosphate of GTP in GTP-bound conformation of EF-Tu are highlighted in orange. (**b**) Atomic Fluctuations of EF-Tu. For each domain of EF-Tu, D1, D2 and D3, the root mean square fluctuation (rmsf) for each residue is shown for each simulation. The P-loop, switch 1 and switch 2 regions of domain D1 are highlighted in grey. Black bars denote the residues that are unresolved in structures of ribosome-bound EF-TU in complex with GDP and KIR [15, 22].

Here, the criterion for a contact between a residue and the *β*-phosphate was that any non-hydrogen atom of the residue was within 5 Å of the *β*-phosphate. The contacts with the P-loop which were present in the GTPase-activated state [3] (residues 19–24) remained present in all the simulations with and without the antibiotic KIR. At the same time, we observed a rapid loss of contacts between GDP and the switch 1 loop in all simulations. In one of the simulations which included KIR (+KIR sim1), we observed spurious occurence of contacts between the *β*-phosphate and residues 60–61 as well as a contact with residue 59, which is not present in the GTP-bound conformation [18]. The loss of contacts between switch 1 and GDP suggests that it is the presence of the *γ*-phosphate that keeps the switch 1 region in place. This result is consistent with the structures of EF-Tu·KIR·GDP·aa-tRNA bound to the ribosome, where GDP is bound to P-loop and the switch 1 region is not resolved [15, 21, 22]. The observation that the switch 1 region is not resolved in these structure indicates that the region is more flexible than the rest of EF-Tu. To check if the loss of contacts between GDP and switch 1 results in an increased flexibility of the switch 1 region, we calculated the root mean square fluctuation (rmsf) of the backbone atoms in all domains of EF-Tu (Fig. 2b). Indeed, the switch 1 region is markedly more flexibile, specifically when compared to the P-loop and the switch 2 region. The residues with large flexibility correspond well with the unresolved residues in KIR-bound structures [15, 22] (Fig. 2b, black horizontal bars).

These results suggest that after P_i_ release, the switch 1 region is not anchored to the rest of domain D1 anymore and can explore a larger range of conformations.

### Interdomain motions of EF-Tu

To elucidate the consequences of the higher switch 1 flexibility on the EF-Tu conformation, we first investigated the interdomain motions of EF-Tu. The two interfaces between domains D1 and D2 and between D1 and D3 are funcionally important. The D1-D2 interface is where the CCA-tail of the tRNA along with the attached amino acid are bound. The observation that deacylated tRNAs have a lower affinity to EF-Tu underscores the importance of this interaction for a stable ternary complex [49, 50]. Further, antibiotics that bind to the D1-D2 interface prevent the binding of a tRNA to EF-Tu and therefore the formation of the ternary complex [51]. The D1-D3 interface is the binding site of the antibiotic KIR which hinders the release of the tRNA from EF-Tu. Notably, the D1-D3 interface is observed to be closed in X-ray structures of the isolated ternary complex with a distance of 6.5 Å between the C*α* atoms of residues Q124 (D1) and R373 (D3) [11]. The interface remains closed when EF-Tu binds to the ribosome in the initial-binding and the codon-recognition states with distances of 5.3 Å and 5.1 Å, respectively [3]. Upon GTPase activation, the interface opens (9.5 Å, [3]) and even more so when KIR is bound (11.6 Å, [22]). The D1-D2 interface remains closed around the CCA-tail of the tRNA in all of these steps.

EF-Tu domains D2 and D3 stay in a very similar conformation relative to each other, both, in all the structures and during the simulations. To investigate the motion of EF-Tu after hydrolysis and P_i_ release, we used principal component analysis (PCA) to extract the dominant conformational modes of motion of D1 relative to D2 and D3 [45]. Figure 3a shows the reaction coordinates of the D1 motion, i.e., the projections onto the dominant mode, as well as the distances between EF-Tu residues H66-E259 and Q124-R373, which report on the closing or opening of the D1-D2 and D1-D3 interfaces, respectively. This D1 reaction coordinate describes a rotation of D1 around the switch 2 helix. To indicate which conformations are observed in the simulations, the logarithm of the probability densities obtained from the simulations are shown for the ribosome-bound ternary complex without and with KIR (with GDP, upper panels) as well as for the isolated ternary complex (with GTP, lower panel). For sufficient sampling, this quantity would represent the free-energy landscape that governs the conformational dynamics in units of *k*_*b*_*T*. The most probable conformations with and without KIR are highlighted with red and green crosses, respectively, and their conformations are shown in Fig. 3b. Residues H66 and E259 interact with the amino acid and the CCA-tail of the tRNA in all states prior to GTP hydrolysis as well as in the KIR-bound GDP state. In the absence of KIR, the D1-D3 interface closed fully in one of the simulations (Fig. 3a, sim1, light green outline) and partially in the other (sim2, dark green outline). In the case of full interface closure, the Q124-R373 distance dropped rapidly during the first 200 ns from the initial 12 Å observed in the KIR-bound cryo-EM structure to around 6 Å and the interface remains closed for the rest of the simulation. This distance is close to the distances observed in the isolated ternary complex as well as the initial-binding and codon-recognition states [11, 3]. In the case of partial closure, after an intial drop to around 9 Å, the distance returns to around 12 Å. The Q124-R373 distance correlates strongly with the D1 reaction coordinate. Notably, the motion of D1 also correlates with an opening of the D1-D2 interface, as can be seen by an increase in the H66-E259 distance to around 16 Å. This extent of D1-D2 interface opening is not seen in structures of EF-Tu with either GTP or with GDP·KIR, where the H66-E259 distance ranges only from 13.3 Å to 14.2 Å [11, 22, 3].

**Figure 3:**
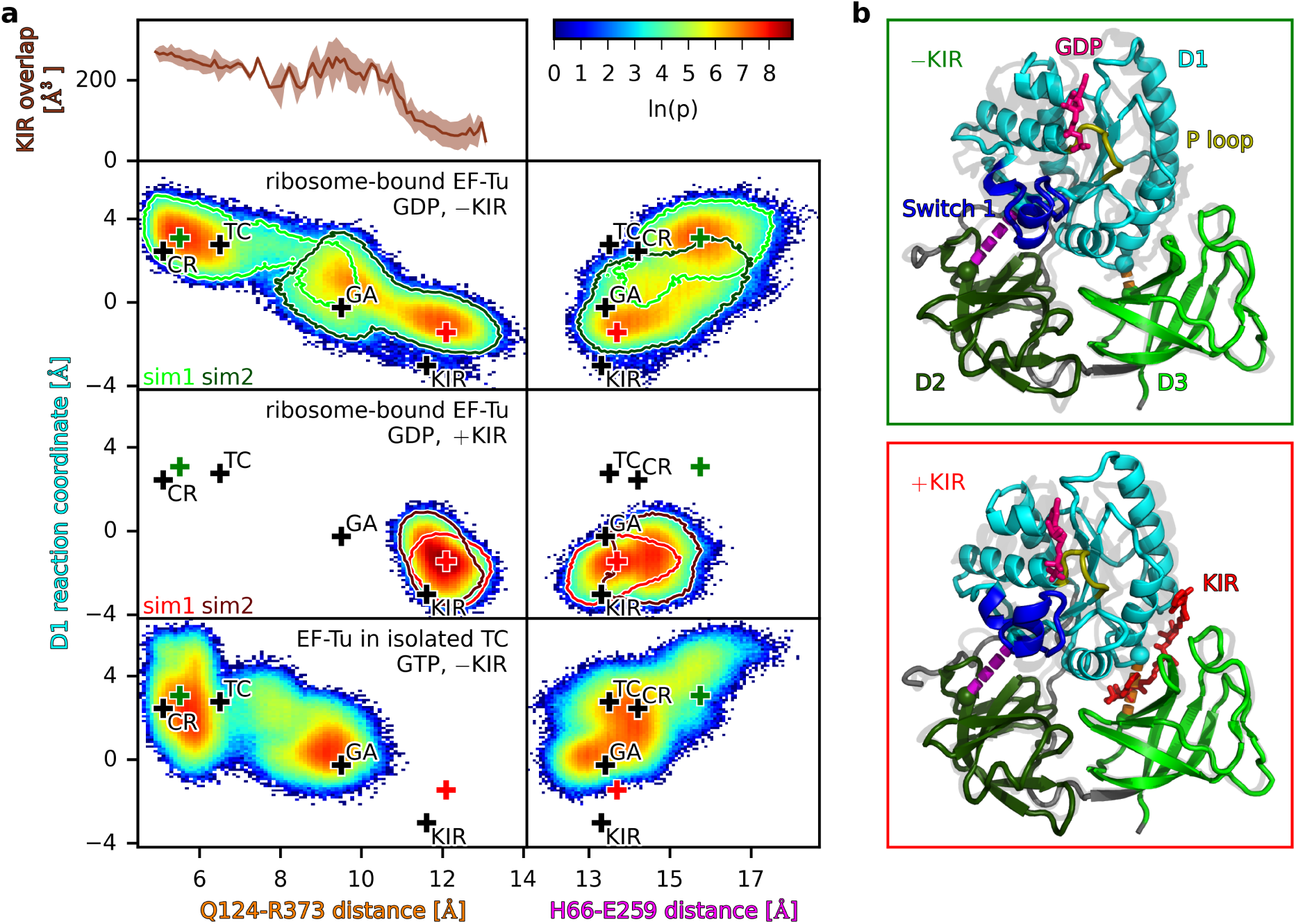
Conformational dynamics of ribosome-bound EF-Tu with GDP with and without KIR, as well as EF-Tu in the ternary complex with GTP. (**a**) The trajectories of the MD simulations were projected onto the reaction coordinates of the D1 motion and the C*α* distances between residues Q123 and R373 as well as H66 and E259. The color represents the logarithm of the probability density *ρ*. Black crosses correspond to the positions of available structures of EF-Tu: in the ternary complex with GTP (TC, [11]), bound to the ribosome in the codon-reading (CR, [3]) and GTPase-activated (GA, [3]) states as well in the KIR-bound state with GDP (KIR, [22]). Red and green crosses indicate the largest probability densities with and without KIR, respectively. For simulations of ribosome-bound EF-Tu, the outlines correspond to ln(*p*) = 2 for simulations sim1 and sim2. KIR overlap as a function of Q124-R373 distance denotes the van-der-Waals overlap between the KIR atoms and EF-Tu atoms after rigid-body fitting KIR into all trajectories of ribosome-bound EF-Tu. The conformations corresponding to the largest probability densities with and without KIR (red and green crosses, respectively) are depicted (upper and lower right panels).

In both simulations with KIR, the D1 reaction coordinate as well as the Q124-R373 and H66-E259 distances remain close to their respective values in the GDP·KIR bound cryo-EM structures [22], which is expected and therefore supports the validity of the starting structure and the simulation setup (Fig. 3a). To investigate if the presence of KIR in the D1-D3 interface sterically prevents the closing, we calculated the van-der-Waals overlap between KIR atoms and EF-Tu after rigid-body fitting KIR atoms into the binding site of structures obtained from the simulations with and without KIR. Fig. 3a shows that this overlap increases with decreasing Q124-R373 distances, indicating that KIR indeed acts as a steric block, preventing the closure of the D1-D3 interface and thereby the opening of the D1-D2 interface.

In summary, the results suggest that upon GTP hydrolysis, P_i_ release results in a loss of the switch 1 loop anchoring to the rest of D1, which frees D1 to rotate around the switch 2 helix. This rotation closes the D1-D3 interface and opens the D2-D3 interface, possibly decreasing the interaction of EF-Tu with the amino acid and the CCA-tail of the tRNA, and therefore the affinity of the tRNA to EF-Tu. The presence of KIR interupts this sequence of events; it thus does not interfere with GTP hydrolysis and P_i_ release, but sterically hinders the closing of the D1-D3 interface which in turn prevents the rotation of D1 and the opening of D1-D2 interface.

### Effect of the ribosome on the D1-D3 interface conformation

The D1-D3 interface of ribosome-bound EF-Tu is in an open conformation in the GTPase-activated state and even further opened the KIR stalled state (Fig. 3a). In all other structures of the ternary complex, whether bound to the ribosome or free, the D1-D3 interface is closed. The spontaneous closure of this interface, observed in our simulations, raises the question if the closed conformation is intrinsically favorable for EF-Tu and the open conformation is induced by interactions with the tRNA and ribosome in the GTPase-activated state. If this were the case, the open interface conformation could resemble the loaded spring which is released only upon GTP hydrolysis and subsequent P_i_ release, as suggested earlier [4]. To test if this is actually the case, we performed six additional simulations of the free ternary complex in solution started from the open GTPase-activated conformation [3].

Figure 3a (lower panel) shows the H66-E259 and Q124-R373 distances relative to the D1 motion for these simulations. In 5 out of the 6 simulations, the Q124-R373 distance decreased to around 6 Å, thereby closing the D1-D3 interface after simulation times ranging from less than 10 ns to 1 *µ*s (Fig. 4a). The fact that EF-Tu transitions to a closed conformation in all but one of the simulations — without back transitions — implies that the closed conformation is indeed energetically favourable in the absence of the ribosome. To further test this conclusion, we calculated the free-energy landscape of the interface closure by additional umbrella sampling simulations. Figure 4b shows the resulting free-energy landscape as a function of the D1-D3 domain closing motion (D3 reaction coordinate, see Methods). Clearly, the closed state is energetically more favorable, which combined with the spontaneous closing in the GDP-bound simulations, further corroborates the idea that the D1-D3 interface opens in the GTPase-activated state by interaction with the ribosome.

**Figure 4:**
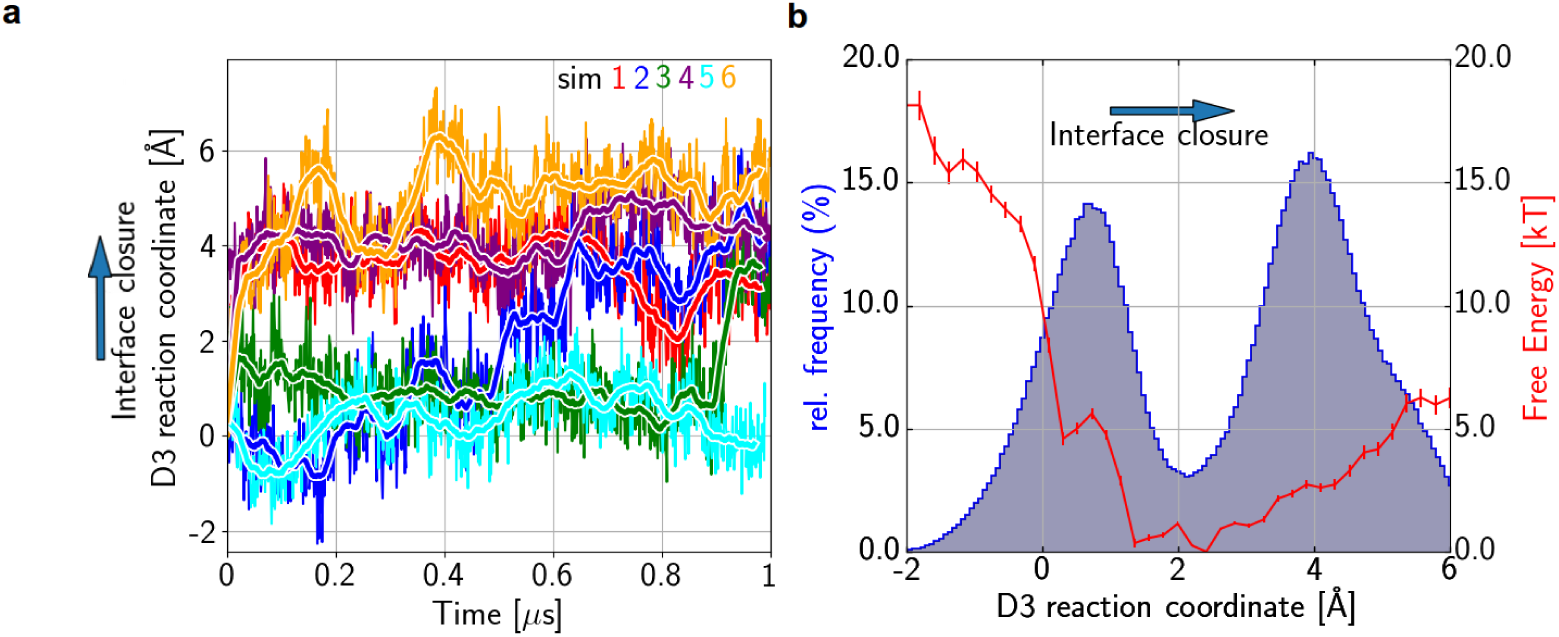
Spontaneous D1-D3 interface closure of EF-Tu in the isolated ternary complex. (**a**) For 6 unrestrained MD simulations, started from a conformation with an open interface, the projection onto the D3 reaction coordinate describing the interface closure is shown as a function of simulation time (colored lines; thick lines: running averages). (**b**) Potential of mean force (PMF, red line) as a function of the reaction coordinate. Error bars were estimated by bootstrapping. The histogram of the concatenated projections of the six unrestrained MD simulations is shown (blue line).

Whereas the D1-D3 interface closed in almost all GTP-bound ternary complex simulations, the D1-D2 interface remained closed around the CCA-tail of the tRNA in all simulations (Fig. 3a). This result, together with the increased flexibility of the switch 1 loop observed in GDP-bound simulations of EF-Tu in the ribosome, provides additional support to the idea that the interaction between the switch 1 loop and the *γ*-phosphate stabilizes the D1-D2 interface in the closed conformation.

### Change of interactions between EF-Tu and aa-tRNA

For the tRNA to fully accomodate into the ribosomal A site, it has to dissocate from EF-Tu. The CCA-tail of the tRNA (nucleotides 74–76) and the attached amino acid (F77) are sandwiched between EF-Tu domains D1 and D2, where A76 and F77 are in contact with E259 and H66 of EF-Tu, respectively [11, 22, 3]. The opening of the D1-D2 interface observed in the simulations of ribosome-bound EF-Tu after removal of KIR suggests that this conformational change may weaken the interaction between EF-Tu and the aa-tRNA. To test this possibility, we calculated the interaction enthalpies of F77 with all D1 and and with all D2 residues (Fig. 5). F77 shows interactions with both D1 and D2 domains in the presence and absence of KIR. In the absence of KIR, when D1 is further away from D2 (increased D1 reaction coordinate), the interaction between F77 and D2 is weakened (less negative interaction enthalpies). Also the interaction of F77 with D1 is weaker on average upon opening of the interface. Closer analysis of the obtained trajectories revealed that when the D1-D2 interface opened, A76 of the tRNA remained bound to domain D2 while F77 tilted away from D2, maintaining weak interactions with D1 (Fig. 5). This conformational change results in a overall weaking of F77 interactions with EF-Tu, which we therefore suggest as the primary step of the tRNA 3’ end dissociation from EF-Tu.

**Figure 5:**
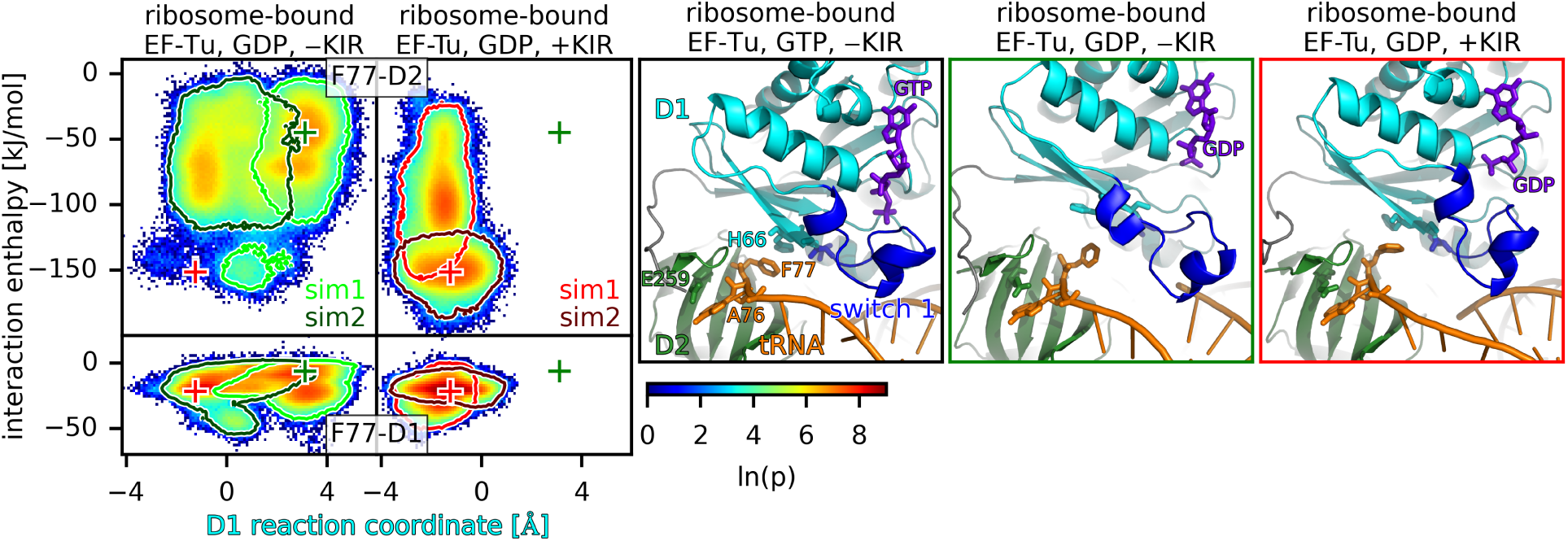
Interaction of ribosome-bound EF-Tu with the amino acid F77 attached to the tRNA. For simulations in the presence and absence of KIR (left and second to left panels, respectively), the probability density is shown as function of the D1 reaction coordinate and the interaction enthalpy of the amino acid attached to the tRNA (F77) with EF-Tu domains D1 and D2 (lower and upper panels, respectively). The green and red crosses denote the positions of the largest probability density obtained from the simulations without and with KIR, respectively. The three rightmost panels show the conformation of the D1-D2 interface obtained from a cryo-EM structure with non-hydrolyzable GTP [3] and from the simulations without and with KIR, corresponding to the green and red crosses, respectively. D1, D2 and the switch 1 loop are colored in cyan, green, and blue, respectively, together with the tRNA (orange) and GTP or GDP (purple).

## Conclusions

Sucessful decoding of the mRNA codon in the ribosomal A site by the ternary complex EF-Tu·GTP·tRNA results in a GTPase-activated conformation of EF-Tu [3]. After GTP hydrolysis and the release of the inorganic phosphate P_i_, the ribsome-bound EF-Tu undergoes a conformational change which preceeds release of EF-Tu from the ribosome [4, 17]. This conformational change ultimately results in the release of the tRNA from EF-Tu, allowing the tRNA to fully accommodate into the A site and EF-Tu to dissociate from the ribosome. Here, using MD simulations started from a cryo-EM structure of the ribosome in complex with EF-Tu·tRNA·GDP which has been stalled by KIR [22], we have investigated the primary conformational changes and energetics of EF-Tu after hydrolysis, and how they might lead to tRNA release.

In structures of ribosome·EF-Tu complexes with inhibited GTP hydrolysis, the switch 1 loop of domain D1 of EF-Tu is resolved and interacts with the tRNA and the *γ*-phosphate of GTP [3, 13]. In our simulations of the state after hydrolysis and P_i_ release, we observed a loss of the contacts between the switch 1 loop and GDP (Fig. 6). GDP remained firmly bound to the rest of the D1 domain, while the switch 1 loop interacted with the tRNA throughout the simulations and was markedly more flexible than most other parts of EF-Tu. This result can explain why in a cryo-EM structures of the ribosome-EF-Tu complex, in which the dissociation after GTP hydrolysis is prevented by the antibiotic KIR, switch 1 loop is not resolved [15, 21, 22]. Vice versa, the fact that the same part that shows high flexibility in our simulations is not resolved in the structures, provides independent support for our simulations. After the loss of the interactions between the switch 1 loop and the rest of D1, which, before hydrolysis, were stabilized by the *γ*-phosphate, D1 is free to rotate towards D3. This rotation results in closing of the D1-D3 and opening of the D1-D2 interface, within which the CCA-tail of the tRNA and its attached amino acid are bound. We therefore propose that, before hydrolysis, the switch 1 loop anchors domain D1 to D2 via the interaction with the *γ*-phosphate. In our simulations, the opening of the D1-D2 interface reduced the interaction between EF-Tu and the amino acid attached to the tRNA, while the tRNA interactions with D3 remain. This result agrees with and explains in structural terms previous ensemble kinetics experiments which indicated that the 3’ end of the tRNA dissociates from EF-Tu first, preceeding the full dissociation [6].

**Figure 6:**
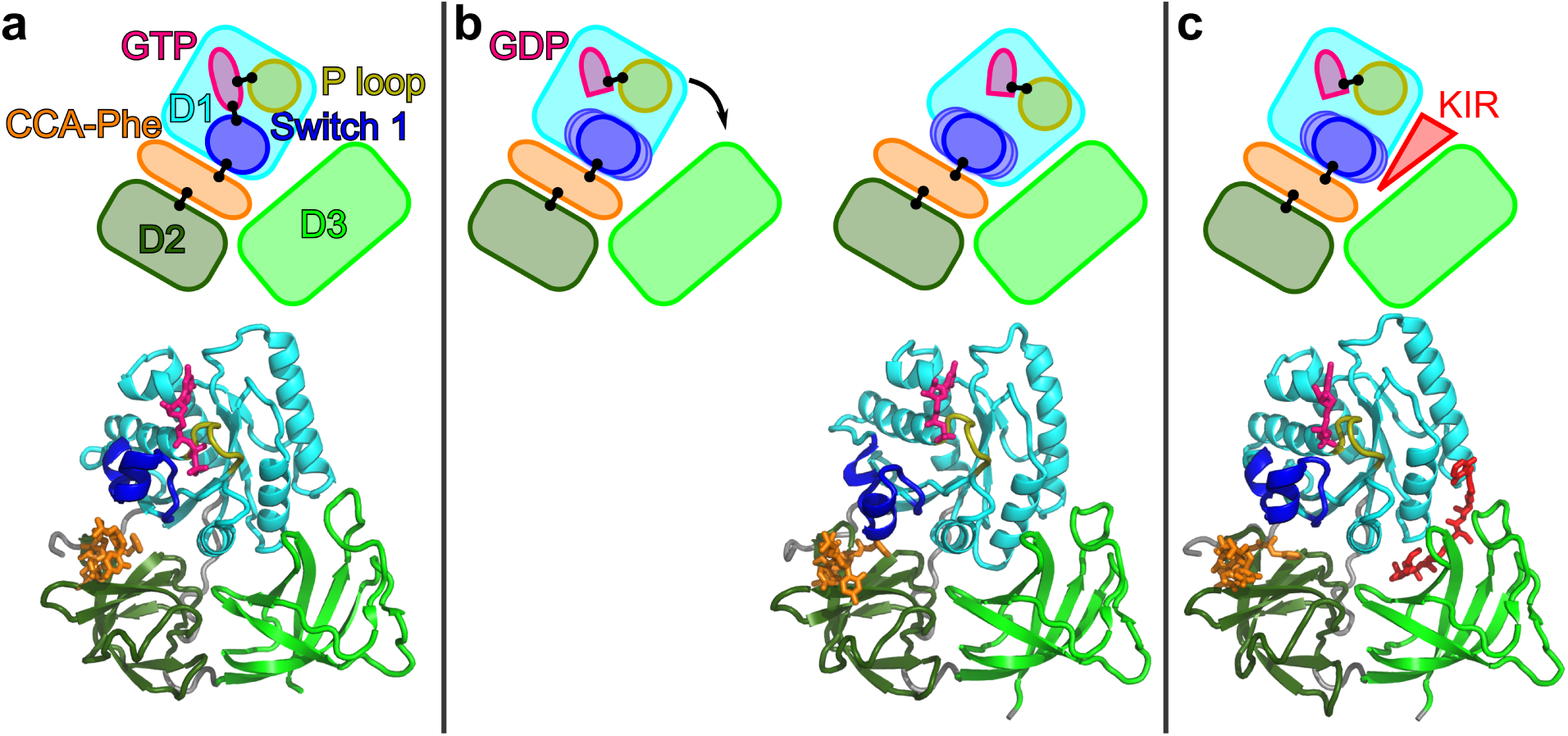
Proposed primary step of EF-Tu after GTP hydrolysis and the mechanism of KIR. (**a**) The conformation of the ribosome-bound EF-TU in the GTPase acivated state. Upper panel: Sketch of the domains of EF-Tu (D1, D2, D3 in cyan, dark green, light green), specific parts of D1 (switch 1 loop, P loop, in blue, yellow) and the tRNA CCA-tail with attached Phe (orange), GTP (pink) and KIR (red). Contacts discussed in the text are represented by black lines. Lower panel: structure of the GTPase-activated state of EF-Tu [3]. (**b**) After hydrolysis to GDP (pink) and P_i_ release (not shown), the contact between GDP and the switch 1 loop is lost, the switch 1 loop becomes flexible. Since D1 is not restrained by the contact to switch 1, it rotates (black arrow) towards D3 into an energetically favourable conformation, thereby opening the D1-D2 and closing the D1-D3 interface. The structure of the last frame of the MD simulation in absence of KIR (sim1) is shown. (**c**) KIR bound to the D1-D3 interface sterically blocks the D1 rotation. The structure of KIR stalled EF-Tu is shown [22].

The weakening of interaction enthalpies between F77 and EF-Tu, upon D1-D2 interface opening, was observed between F77 and D2 residues and, to a lesser degree, between F77 and D1 residues. The interaction between F77 and D1 was mainly with histidine 66, which strongly contributes to the affinity of Phe-tRNA^Phe^ to EF-Tu and to a lesser degree (or not at all) to that of other tRNAs [52]. This specificity of the interaction between the attached amino acid and EF-Tu suggests that the details of dissociation of the 3’ end of the tRNA most likely depend on the amino acid. Nevertheless, we would expect the opening of the D1-D2 interface to destabilize the interaction between EF-Tu and all other amino acids as well.

Due to the relatively short timescales that can currently be covered with unrestrained explicit-solvent all-atom MD simulation [53], the subsequent steps of full EF-Tu dissociation from the ribosome and the tRNA are not observed in our simulations. Based on biased MD simulations, Lai et al. proposed a pathway from the closed GTP-bound conformation to the open GDP-bound conformation which involves a separation of D1 from D2 and D3, followed by a rotation of D1 and a subsequent rejoining of D1 with D2 and D3 in the GTP-bound conformation [54]. Unrestrained coarse-grained MD simulations of the transition of EF-Tu between the GTP- and the GDP-bound conformation conformation suggested that multiple routes exist, a direct path, paths involving domain separation, and more disordered intermediate conformations [55]. A combination with coarse-grained simulations of tRNA accomodation [56] suggested that these disordered conformations affect the kinetics of tRNA accommodation [55]. In agreement with our observations, no spontaneous transitions were observed in unbiased all-atom MD simulations on a *µ*s-timescale [54, 55].

Interestingly, the D1-D3 interface is closed in all structures of the isolated GTP-bound ternary complex and in ribosome-bound conformations, except for the GTPase-activated structure [3]. This observation indicates that the EF-Tu conformation with an open D1-D3 interface is stabilized by interactions with the ribosome in the GTPase-activated state. It also suggests that upon GTP hydrolysis, D1 is able to rotate and reverts back into the energetically preferred closed D1-D3 conformation.

The antibiotic KIR binds to the D1-D3 interface and thereby sterically hinders the D1 rotation. As a result, the D1-D2 interface remains closed around the 3’ end of the tRNA in our simulations including KIR, maintaining strong interactions with the 3’ end, thereby preventing the dissocation of the 3’ end from EF-Tu. We propose that the prevention of this first dissociation step prevents full EF-Tu dissociating from the tRNA and the ribosome after GTP hydrolysis and thus locks EF-Tu into a conformation close to the pre-hydrolysis conformation [19, 20].

Several other antibiotics are similar to KIR in structure and binding site, and therefore the proposed mechanism may apply to a larger class of antibiotics [57, 58, 59, 60]. For example, aurodox has been shown to occupy the same binding site in the same conformation as KIR [60]. Further, this structure contains GDP and the switch 1 loop is not fully resolved, indicating that aurodox stalls EF-Tu on the ribsome in the same way as KIR. But also structurally unrelated antibiotics, such as Enacyloxin IIa, bind to the D1-D3 interface in a binding site which overlaps with that of KIR [61, 62], result in an open D1-D3 interface, and prevent the dissocation of EF-Tu·GDP from the ribosome. The observation that a differently structured antibiotic, which occupies the same binding site as KIR, has the same effect, is further experimental support for the idea that KIR sterically hinders the domain closure, which appears to be a necessary primary step of EF-Tu release from the tRNA and the ribosome, and suggests a general mechanism.

## Acknowledgement

This research was supported by the DFG Forschergruppe FOR1805 (to H.G. and L.V.B.). We thank the computer center Garching (RZG) and the Gesellschaft für wissenschaftliche Datenverarbeitung Göttingen (GWDG) for technical assistance; computer time has been provided by the RZG.

